# Phase separated liquid vimentin droplets stabilize actin fibers through wetting

**DOI:** 10.1101/2024.06.15.597620

**Authors:** Arkaprabha Basu, Tommy Krug, Benjamin du Pont, Qiaoling Huang, Sijie Sun, Stephen A. Adam, Robert Goldman, David A. Weitz

## Abstract

The cytoskeleton is composed of F-actin, microtubules, and intermediate filaments (IFs). Vimentin is the most ubiquitous IF. It is involved in wound healing, tissue fibrosis and cancer metastasis, all of which require rapid vimentin filaments assembly. In this paper, we report that un-polymerized vimentin forms liquid condensates that appear to enable rapid filament growth. Given the transient nature of these droplets, we focus on properties of vimentin-Y117L, a mutant which does not form filaments, enabling us to study these droplets in detail. They dissolve under 1,6-Hexanediol treatment and under decreasing concentration, confirming that they are liquid, and phase separated. These condensates extensively wet actin fibers, rendering them resistant to actin-depolymerizing drugs. We show similar behavior occurs in wild type vimentin during its assembly into filaments.

## Introduction

The cytoskeleton is a set of biopolymer networks that is responsible for the motion and mechanical properties of the cell, as well as intracellular cargo transport^1^. In eukaryotic cells, it is primarily made of three interpenetrating networks: actin^2^, microtubules^3^, and intermediate filaments^4^. The first of these networks is the actin cytoskeleton, which is a network formed by actin filaments and actin-binding proteins such as myosin, *α*-actinin and filamin. At the edge of the cell, densely packed actin filaments, the cortex, give the cell its mechanical properties. Actin also forms stress fibers, thick bundles of cross-linked actin filaments, which are responsible for cellular motion, contractility and mechanotransduction. For example, actin filaments cross-linked by myosin motor proteins, are the main contractile machinery of the cell. The second network, microtubules, is the primary instrument for intra-cellular cargo transport. Structurally, they are hollow tube-like polymers of alternating alpha and beta tubulin sub-units. Along microtubules, motor proteins like kinesis and dynein transport vesicles and organelles throughout the cell. During cell division, microtubules form the mitotic spindle, which is responsible for chromatin separation. The third component of the cytoskeleton, intermediate filaments (IFs), is more varied; it encompasses a host of proteins, including vimentin^5,6^, neurofilaments^7^ and keratins^8^. Vimentin, the most ubiquitous intermediate filament, is responsible for would healing in healthy tissues. The vimentin network also plays a key role in cancer cells undergoing metastasis; it provides the cell with the unique mechanical properties necessary to squeeze through the dense tumor tissue environment.

These networks are highly dynamic, and their assembly and disassembly are involved in important cellular processes. For example, actin polymerization at the leading edge of the cell, coupled with depolymerization on the opposite side, drives cellular movement in keratinocytes^9^. In microtubules, the most drastic dynamics occurs during mitosis where the entire microtubule network dissolves and reassembles into mitotic spindles. The vimentin network breaks down during mitosis, forming small punctate structures which reorganize into filaments in the daughter cells. In all of these instances, the filaments grow within minutes. This cannot be explained by randomly diffusing sub-units organizing into polymers. Though sub-units diffuse through the cytoplasm quickly because of their low molecular mass, they must have the correct orientation to be incorporated into the filaments, greatly slowing filament elongation.

A possible mechanism for cells to increase rates of assembly of filamentous structures is by concentrating sub-units and related proteins locally, with the filament growth occurring inside these regions. For example, actin-associated proteins, NcK, N-WASP and Nephrin, form phase separated condensates with increased concentrations of actin monomers and an actin polymerizing protein, Arp2/3^10^. This enables a 14-fold increase of actin polymerization rate. Similarly, rapid growth of microtubules is enabled by centrosomes, which are phase separated condensates, enriched with tubulins and their assembly-related proteins such as gamma-tubulin ring complex proteins and XMAP215^11^. By contrast, vimentin assembly does not require any other proteins. Instead, it is well established that vimentin assembly is a multi-step process; it forms tetramers which laterally assemble into unit length filaments (ULFs) and finally elongate into vimentin intermediate filaments (VIFs)^12–14^. However, it is unclear how VIF growth can be so rapid in the absence of any other proteins, which can increase the vimentin concentration.

In this paper, we shown that vimentin, by itself, forms phase separated condensate state, which is a precursor to VIF assembly, and can thus account for its high polymerization rates. We focus on the mutant vimentin-Y117L^15–17^, which cannot form VIFs, allowing us to study the properties of the condensates. They have many hallmarks of liquid droplets: they merge, split, undergo shape fluctuations, and selectively dissolve under 1,6-hexanediol treatment. Moreover, their rapid and reversible dissolution with decreasing concentrations is consistent with them being formed through liquid-liquid phase separation (LLPS). Unlike vimentin filaments which mostly interact with microtubules^18,19^, these condensates primarily interact with actin fibers. A large fraction of these droplets can wet actin fibers and remain attached to them for a long time. Such fiber-attached droplets exhibit a specific motion: it is diffusive at short time-scales but directional along the fibers at longer time-scales. This directed, or ballistic, motion is not driven by motors but originates from the retrograde flow of actin. When the actin fibers are wet by the vimentin droplets, they become unrecognizable to actin-binding molecules. This, in turn, protects them from depolymerizing drugs like Cytochalasin B. We also demonstrate that wild type vimentin, during assembly, goes through similar phase separated liquid condensates before forming VIFs. These results suggest that local increases in concentration due to phase separation may lead to more rapid assembly of vimentin intermediate filaments.

## Results and Discussion

### Vimentin forms liquid droplets through phase separation

The network formed by vimentin intermediate filaments (VIFs) is an integral part of the cytoskeleton which determines the mechanical properties of a cell^5,6^. Once formed, the VIF network incorporates virtually all available vimentin monomers, leaving very few of them free^20^. Interestingly, vimentin can also exist in a punctate state that is distinct from VIFs. For example, when vimentin expression is induced in vimentin-null MCF-7 cells, filamentous networks do not form when its concentration is low. Instead, it first forms small spherical puncta which ultimately transform into the VIF network^21^. Similarly, when Baby Hamster Kidney (BHK) cells undergo mitosis, their vimentin filaments dissolve into small spherical puncta which resemble those in MCF-7 cells and transform back into a VIF network structure in each daughter cell^22^. However, although these punctate states of vimentin are each precursor to the filaments, they are transient; thus, it is not possible to study their properties.

To overcome this difficulty, we use vimentin-Y117L, where the substitution of the 117^th^ amino acid of vimentin from Tyrosine (Y) to Leucine (L) prevents VIF formation; this lets us study the behavior of the puncta in more detail^15^. To avoid interference from other intermediate filament networks we use a cell line derived from Mouse Embryonic Fibroblast (MEF) cells which are devoid of other IFs. This cell line is produced by stably transfecting vimentin knock-out MEF cells with a plasmid containing vimentin-Y117L fused to a fluorescent mEmerald so that the only vimentin is the cell is the fluorescent mutant. By imaging the cells with a confocal microscope, we find their cytoplasm is replete with vimentin puncta that are predominantly spherical, as shown on green in Fig. 1a and Supplementary Video 1. Observing this behavior at higher magnification reveals that the puncta are highly dynamic, undergoing constant shape fluctuations. This can be seen from the different shapes of the same punctum marked by the yellow arrow in Figs. 1b-e, and more clearly in Supplementary Video 2. This behavior is reminiscent of liquid droplets. Even more convincing evidence of such liquid droplet-like behavior comes from observing two spherical puncta that come in contact with one another and merge into a single, larger sphere over 13 seconds as shown by the magenta circle in Figs. 1b-e. The merged puncta remain a single structure, thus eliminating the possibility that they are just random particles overlapping with one another. Thus, these puncta must be liquid droplets and not solid aggregates or membrane-bound organelles.

**Figure 1.**
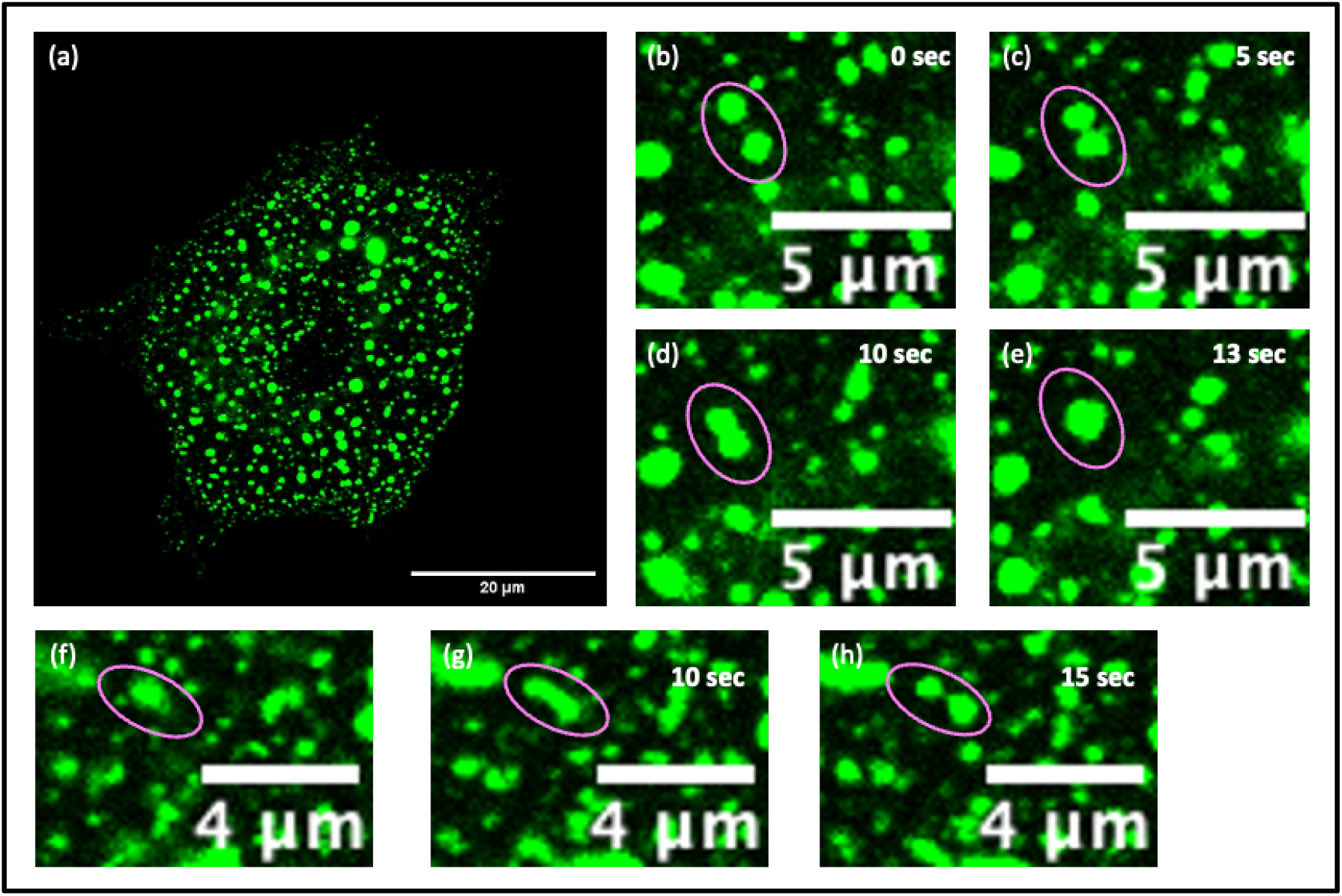
Appearance and dynamics of Vimentin-Y117L puncta. (a) MEF cell containing Vimentin-Y117L-mEmerald shows fluorescent puncta throughout the cytoplasm. (b)-(e) Two spherical Vimentin-Y117L puncta, shown inside the magenta circle, come in contact with one another and merge into a larger spherical puncta over 13 seconds. (f)-(h) A Vimentin-Y117L puncta undergoes shape fluctuation into an ellipse which eventually splits into two smaller spherical puncta.

These liquid droplets also exhibit a very surprising feature: they can attain a highly elliptical shape which can split into two smaller spherical droplets, as illustrated in the magenta circle in Fig. 1f-h. The highly elliptical shape, which is essential for splitting, is unlikely to be due to shape fluctuations alone; it may instead result from shear forces, although these are not directly observed.

To explore whether the vimentin-Y117L droplets are truly liquid, we investigate their response to 1,6-Hexanediol, an aliphatic diol which selectively dissolves membraneless liquid compartments in cells^23,24^. After 2 minutes in 0.5% (w/v) 1,6-Hexanediol cells lose a majority of the droplets, as shown in Fig. 2a. By contrast, when we fix untreated cells, we observe an abundance of droplets, as shown in Fig. 2b. The 1,6-Hexanediol-induced dissolution of the droplets confirms that they are liquid and membraneless. As a control, we compare the effect of the drug on another cytoskeletal structure. We choose actin fibers, which are solid, and treat the cells with the actin-binding dye SiR-actin for visualization. The drug treatment has no obvious effect on the actin network of the cells, which can be seen by comparing actin fibers of drug-treated and untreated cells shown in magenta in Figs. 2c and 2d respectively. This demonstrates that 1,6-Hexanediol does not affect the non-liquid polymeric actin. In contrast, the drug treatment leads to a 10-fold decrease in the average number of droplets per cell, as shown in Fig. 2e.

**Figure 2.**
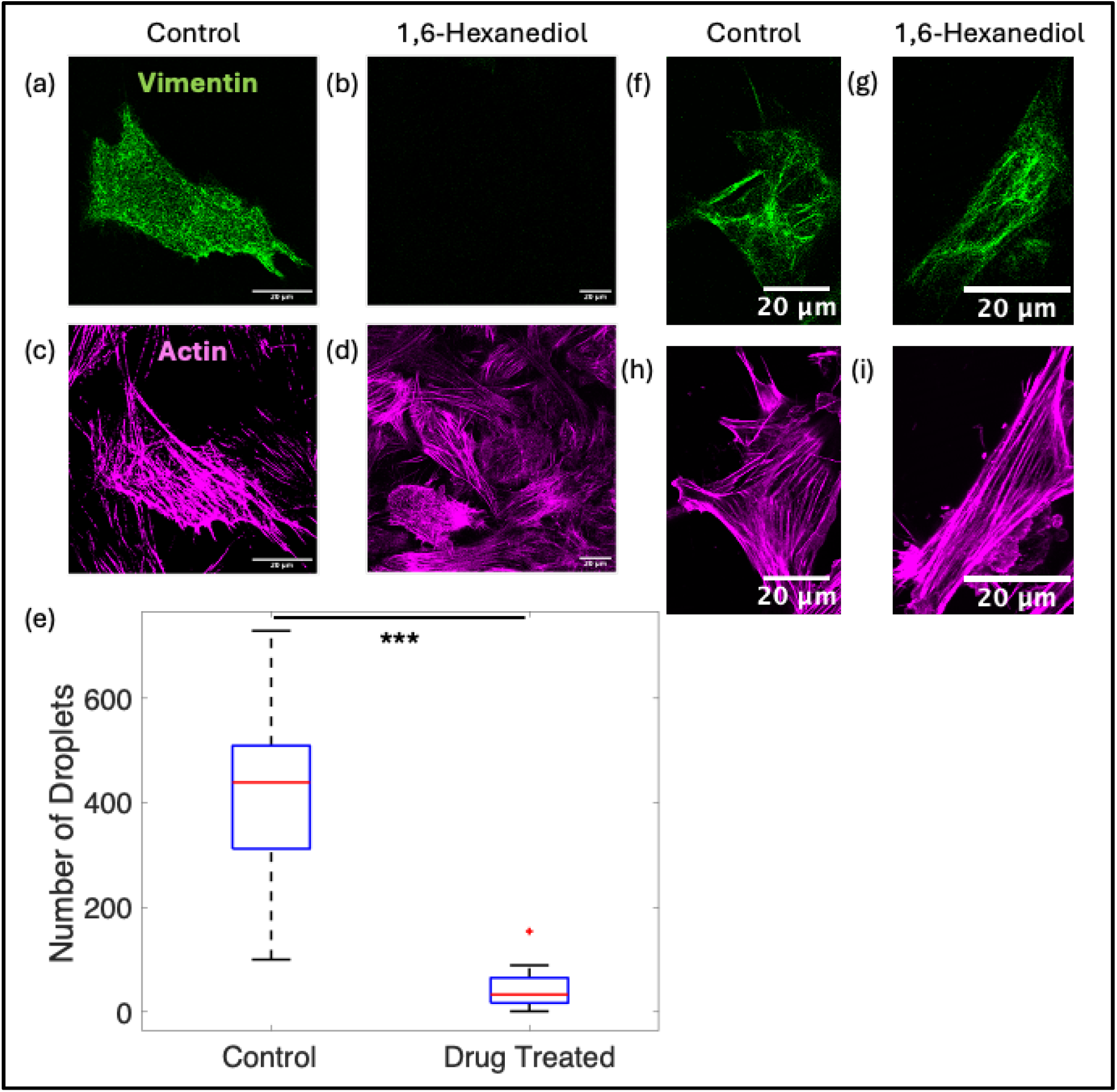
Effect of 1,6-Hexanediol on MEF cells. (a),(c) Vimentin-Y117L droplets (green) and actin fibers (magenta) in untreated MEF cells. (b),(d) Vimentin-Y117L (green) and actin fibers (magenta) in MEF cells after 0.5% 1,6-Hexanediol treatment for 2 minutes showing drastic decrease in Vimentin droplets while actin remains unaffected. (e) Box plot of number of droplets of cells before (N=17) and after (N=22) drug treatment demonstrating statistically significant decrease in number of droplets. (f),(h) Vimentin filaments (green) and actin fibers (magenta) in untreated MEF cells. (g),(i) Vimentin filaments (green) and actin fibers (magenta) in MEF cells after 0.5% 1,6-Hexanediol treatment for 2 minutes showing both actin and Vimentin remain unaffected.

Since, Fusion of vimentin-Y117L with the fluorescent protein mEmerald could affect its physical properties, including the state of the punctate structure, we study cells with non-fluorescent vimentin-Y117L using immunostaining. We again observe abundant droplet-like vimentin structures, as seen in Supplementary Fig. 1a. These droplets again disappear when the cells are treated with 0.5% 1,6-Hexanediol for 2 minutes, as seen in Supplementary Fig. 1b. Thus, the liquid nature of the vimentin-Y117L droplets is an inherent property of the protein and not a result of mEmerald fusion. As a control we again use actin and visualize it by Phalloidin staining. The actin fibers remain unaffected by the 1,6-Hexanediol treatment, as shown in Supplementary Figs. 1c,d.

To test whether the 1,6-Hexanediol-induced dissolution of droplets leads to irreversible alterations of vimentin, we treat the cells with 0.5%-1,6-Hexanediol for minutes, remove the drug, replace with normal culture media to let the droplets reform and visualize actin and vimentin using immunostaining. Vimentin-Y117L droplets reappear within 4 hours of the drug withdrawal, with some cells regaining droplets in under 2 hours as in shown in Supplementary Fig. 2. This confirms that the 1,6-Hexanediol-induced dissolution of vimentin-Y117L droplets is reversible.

To explore the effect of the drug on fully polymerized VIFs, we carry out the same 1,6-Hexanediol treatment on MEF cells which contain vimentin tagged with mEmerald, as shown in Fig. 2f. The drug treatment has no visible effect on the VIFs of these cells, as can be seen by comparing cells with and without drug treatment, where the VIF networks are shown in green in Figs. 2g and 2f respectively. As further control, actin fibers, visualized using SiR-actin, also remain unaffected by the drug treatment; this can be seen by comparing cells with and without drug treatment, where the actin network is shown in magenta, in Figs. 2i and 2h respectively.

The fact that VIFs do not dissolve under conditions where the vimentin droplets do dissolve confirms that the filamentous state of wild type vimentin is distinct from the liquid droplet state of vimentin-Y117L. One way for cells to form membraneless liquid compartments, such as these droplets, is liquid-liquid phase separation (LLPS)^25,26^. This process is driven by weak interactions, causing components of a solution to spontaneously de-mix. The condensates formed through LLPS are typically liquid, owing to the weakness of the underlying interactions responsible for them. Furthermore, intrinsically disordered regions (IDRs) of proteins are often implicated in the weak interactions which give rise to LLPS^27,28^. Vimentin has two terminal IDRs which is consistent with its ability to phase separate^29,30^.

To test whether the droplet formation is indeed driven by LLPS, we explore the phase behavior of vimentin-Y117L. Because phase separation is concentration dependent, we change the concentration of all proteins in the cell by exposing it to culture medium diluted to 20% v/v in water. As the osmotic pressure outside the cell drops, its volume increases due to the influx of water thereby decreasing the concentrations of all cytoplasmic proteins. The extra water in the cell also hydrates the IDRs which alters their interaction, further affecting structures that are already reliant on weak interactions^31^. When cells with abundant vimentin droplets are exposed to diluted medium for 30 seconds, the droplets begin to dissolve, as can be seen by comparing the images of the cell in Figs. 3a and 3b. Within 2 minutes, all droplets disappear and a uniform vimentin fluorescence is observed throughout the cytoplasm, as Fig. 3c demonstrates. Since a change in the concentration and interaction strength should only affect phase separated structures, we also tested the impact of dilution on actin fibers, which are not phase separated and have no IDRs, as a control. In cells treated with SiR-actin, actin fibers are detected throughout the diluted medium treatment. In the time it takes for the vimentin droplets to dissolve entirely, the actin fibers remain unaffected, as can be seen by comparing the actin network of the cell, shown in magenta, in Figs. 3a-c. If this dissolution reflects the phase behavior, it should be reversible. To test this, we remove the diluted medium and add fresh normal medium to the cell and observe both vimentin and actin in the cell. Within 30 seconds, the uniform fluorescent background becomes non-uniform and within 2 minutes droplets start reappearing throughout the cell, as shown in green in Figs. 3d and 3e respectively. The droplets are completely recovered by 5 minutes, as the green in Fig. 3f exemplifies. The actin, however, remains unaffected, as seen in magenta in Figs. 3d-f. These results demonstrate that vimentin droplet formation is due to phase separation.

**Figure 3.**
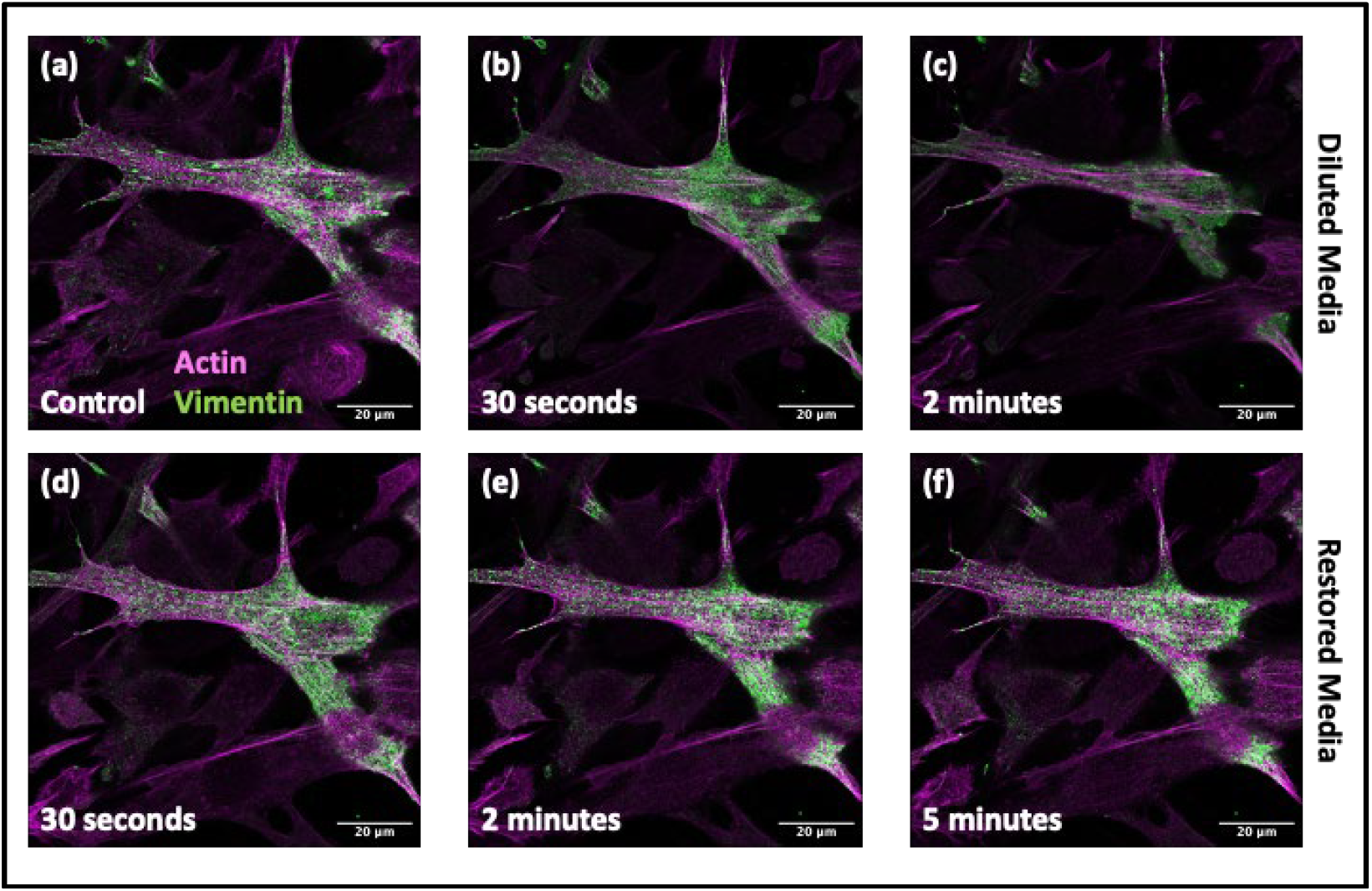
Effect of concentration change on vimentin-Y117L droplets in MEF cells. (a-c) Vimentin-Y117L droplets (green) and actin fibers (magenta) in a cell before (a), 30 seconds after (b) and 2 minutes after (c) diluted media treatment. (d-f) Vimentin-Y117L droplets (green) and actin fibers (magenta) in the same cell after 30 seconds (d), 2 minutes (e) and 5 minutes (f) of restoring fresh media.

### Vimentin droplets colocalize with actin fibers

In our observation of droplets in cells expressing vimentin-Y117L, we notice that they frequently appear in linear arrays reminiscent of the distribution of actin fibers. To explore if vimentin droplets interact with actin, we examine how they are spatially distributed in the cell. A large fraction of vimentin droplets overlap with the actin fibers, as seen by comparing the actin network shown in magenta with the vimentin droplets shown in green in Figs. 4a and 4d respectively. To better visualize this phenomenon, we employ a threshold function to obtain an image, where the areas with overlapping fluorescence signals are white; the extent of colocalization can be seen from the abundance of white areas in Fig. 4c. On average, ∼65% of vimentin is colocalized, as seen in Fig. 4g.

**Figure 4.**
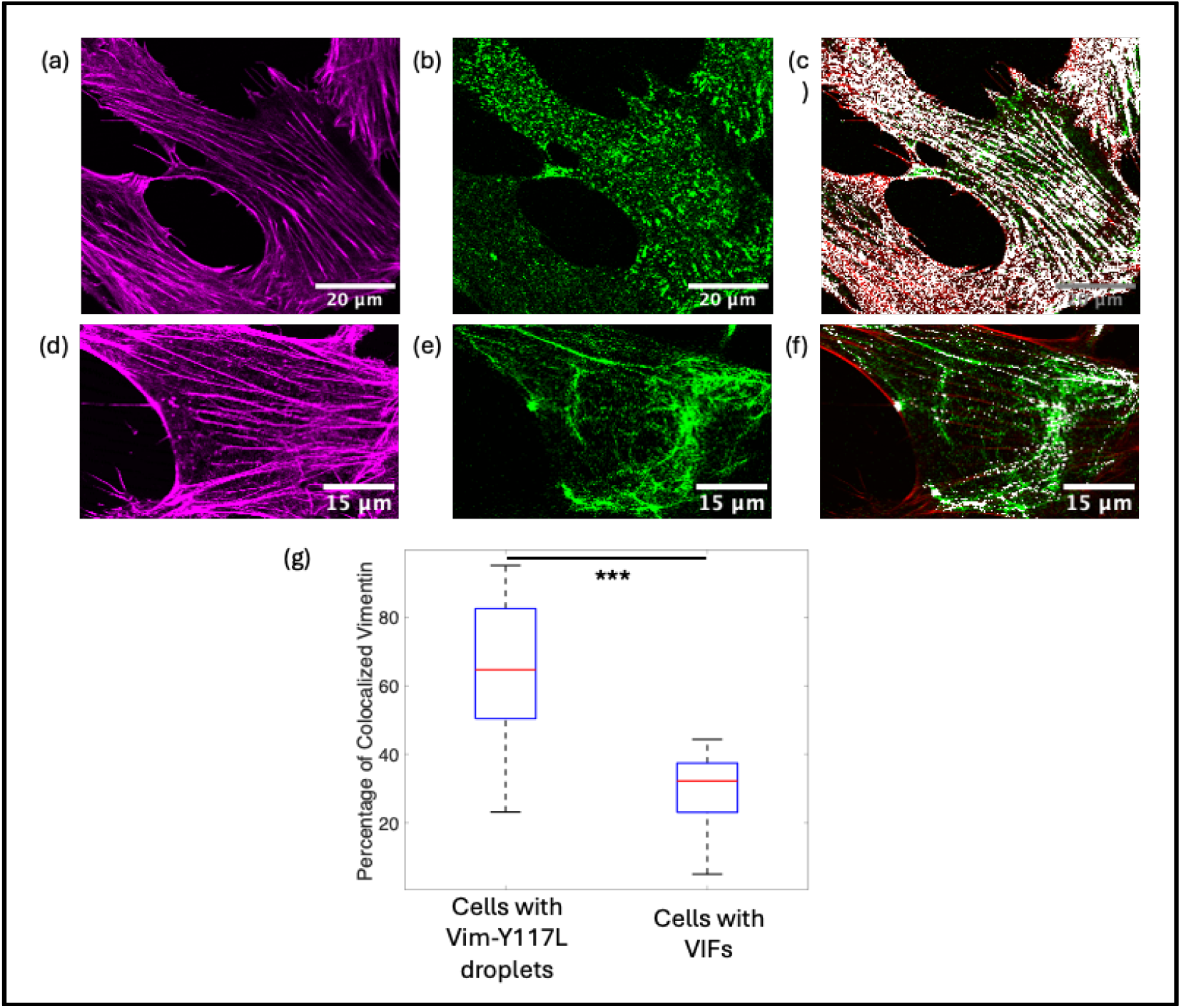
Colocalization between actin and Vimentin. (a,b) Actin (magenta) and Vimentin (green) of a MEF cell with Vimentin-Y117L droplets. (c) Colocalization map of the cell showing colocalized (white) region, un-colocalized actin (red) and un-colocalized Vimentin (green). (c,d) Actin (magenta) and Vimentin (green) of a MEF cell with Vimentin filaments. (f) Colocalization map of the cell showing colocalized (white) region, un-colocalized actin (red) and un-colocalized Vimentin (green). (g) Box-plot of percentage of colocalized Vimentin for MEF cells with VIFs (N=24) and MEF cells with Vimentin-Y117L droplets (N=31).

To study how this behavior compared to cells with vimentin filaments, we quantify the overlap between actin and vimentin filaments. The filament networks are spatially distinct, as seen by comparing the actin network shown in magenta and the vimentin filament network shown in green in Fig. 4d and 4e respectively. The thresholded image for this cell shows mostly independent actin and vimentin signals, as seen by the absence of white areas in Fig. 4f. The average colocalization of VIFs is less than 30%, in stark contrast to the 65% for the droplets, as seen in Fig. 4g. The 30% overlap between VIFs and actin is consistent with recent results which show that actin and vimentin networks are often inter-penetrated in the cell^32^. However, our results demonstrate that the actin-vimentin colocalization behavior is enhanced in the droplet state of vimentin.

To investigate the nature of the interaction between the vimentin droplets and actin fibers, we study the motile properties of the droplets. While free droplets move rapidly in every direction, colocalized droplets move along the fibers they are on, rarely coming off, as seen in Supplementary Video 4. In one example, a droplet travels ∼10 *μ*m along a single actin fiber for ∼9 minutes, as shown by the yellow circles in Figs. 5a-c. Thus, the interaction between the droplets and the actin fibers must be strong enough to prevent detachment. Both these motions are distinct from the microtubule dependent bidirectional motion of these structures^16^. Furthermore, our results demonstrate that the droplets on actin fibers are not static but move in a specific manner.

**Figure 5.**
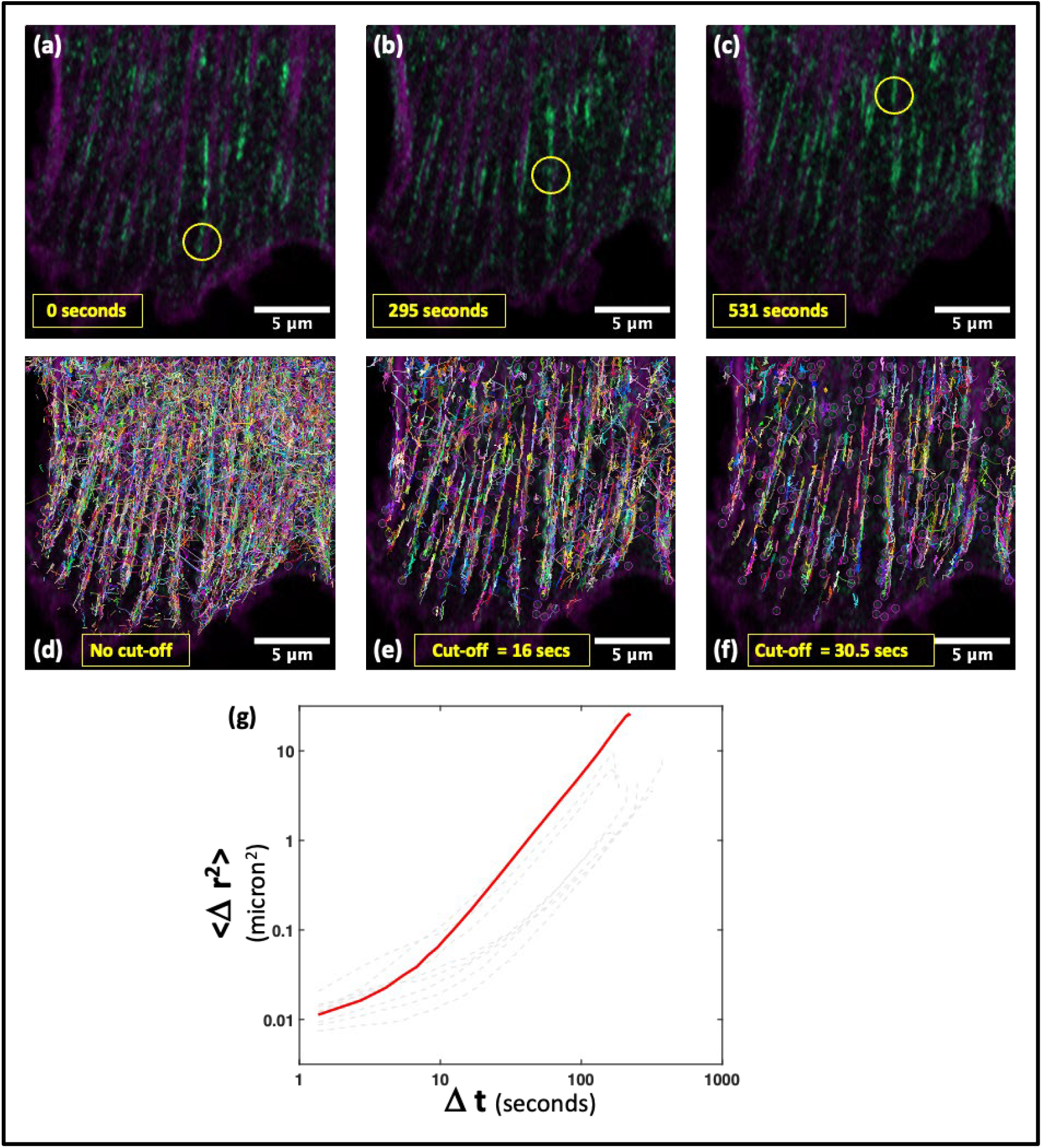
Motion of Vimentin-Y117L droplets. (a-c) A Vimentin-Y117L droplet (green), shown inside the yellow circle traveling ∼ 10 mm along an actin fiber (magenta) over ∼9 minutes. (d) Tracks of all the identifiable Vimentin droplets in the cell overlayed with actin fibers (magenta). (e) Droplet tracks longer than 16 seconds overlayed with actin fibers (magenta). (f) Droplets tracks longer than 30.5 seconds overlayed with actin fibers (magenta). (g) Plot of mean squared displacement, ⟨Δ*r*^2^(*t*)⟩, as a function of time, Δ*t*, for one droplet (red) showing two different slopes. Same plot for eight more droplets shown in dotted grey.

To quantify the motion of the droplets, we determine their trajectories as a function of time. When these are all plotted on the same image, the cell is filled with paths going in all directions; they are jumbled and cannot be visually distinguished from one another, as shown in Fig. 5d. However, longer trajectories predominantly lie along actin fibers. This can be illustrated by plotting trajectories longer than 16 seconds which preferentially eliminates the off-actin movement, as shown in Fig. 5e; doing the same for trajectories longer than 30.5 seconds almost exclusively shows the motile droplets along actin fibers, as shown in Fig. 5f. Being stuck to the actin makes the colocalized droplets stay within the imaging volume longer than free ones, resulting in longer trajectories.

We further investigate the droplet dynamics by measuring their mean squared displacement, ⟨Δr^2^(*t*)⟩, as a function of time difference, Δ*t*. The longest tracks contain both long time-scale and short time-scale behavior; therefore, we use them for this analysis. The plot of ⟨Δ*r*^2^(*t*)⟩ has two distinct slopes, corresponding to two distinct modes of motion, as can be seen from the data shown in the solid curve in the logarithmic plot in Fig. 5g. At short time-scales, the slope is ∼1, indicating the motion Is diffusive with a diffusion coefficient of 2.5 (±1) *μ*m^2^s^-1^. The slope changes to ∼2 for larger time-scales, indicating the motion becomes ballistic, where the droplet moves in a directed manner. The cross-over from diffusive to ballistic motion takes place at Δ*t* ≈7.5seconds. This same diffusive to ballistic transition is replicated by virtually all droplets, as can be seen, for example, from the dashed lines corresponding to other particles, as shown in Fig. 5g. The average velocity of the directed motion in 1.25 (±0.5) *μ*m/minute. This is too low to be driven by motors^33,34^, which move almost ten times faster. Instead, it is consistent with actin retrograde flow^35,36^. This suggests the directional on-actin motion of the droplets result from actin treadmilling, the natural polymerization and depolymerization of actin.

### Vimentin droplets wet and coat actin fibers and protects them

Unlike spherical free droplets, the ones colocalized on actin fibers are often elongated, as shown by the yellow arrows in Figs. 6a and 6b. They can also undergo drastic changes in shape as they move along the actin fibers, as seen by comparing the two shapes of the same droplet shown inside the yellow circles in Figs. 5b and 5c as well as by their expanded images shown in Figs. 6c and 6d. The ability of the droplets to change shape suggests that they remain liquid even when bound to actin fibers. One mechanism for liquid droplets to attach to other structures is wetting, which is a physical phenomenon. When a liquid droplet comes in contact with another structure, they interact with one another only at the surface, or interface, and this surface interaction can often determine their behavior. Wetting is when droplets spread out over the new structure thereby enhancing the interface between them, which can result in the loss of their spherical shapes^37–39^.

**Figure 6.**
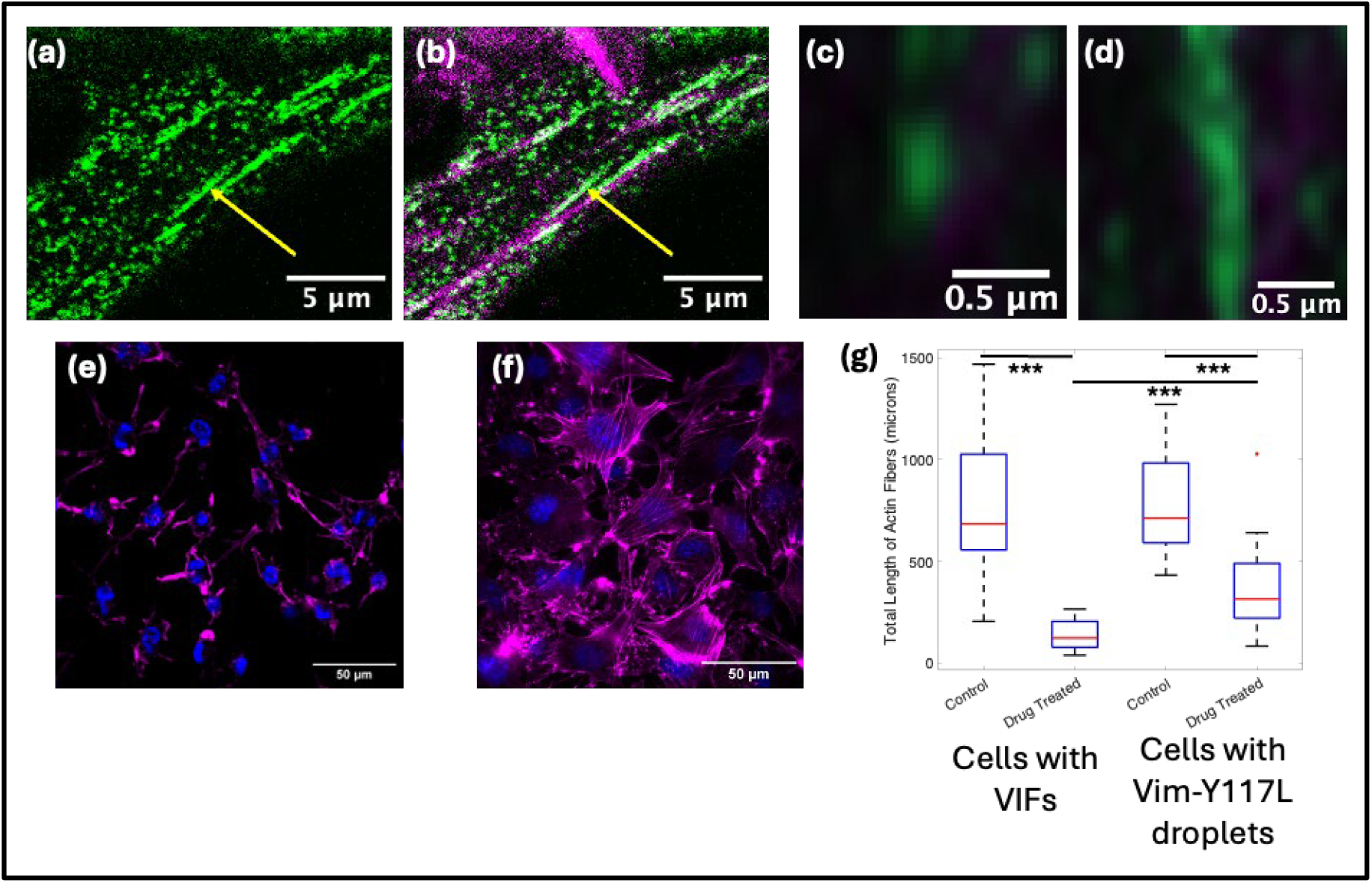
Vimentin droplets coat the actin fibers in MEF cells. (a) Vimentin-Y117L droplets (green) showing elongated droplets with the yellow arrow. (b) Overlay image of the vimentin-Y117L droplets (green) and the actin fibers (magenta) of the same cell; yellow arrow showing that the elongated droplets colocalize with the actin fibers. (c),(d) Expanded images of a vimentin-Y117L droplet traveling along acting fibers exhibiting different shapes. (e) MEF cells with vimentin intermediate filaments showing their nucleus (blue) and actin (magenta) after Cytochalasin B treatment demonstrating almost no cells have actin fibers left. (f) MEF cells with vimentin-Y117L droplets showing their nucleus (blue) and actin (magenta) after Cytochalasin B treatment demonstrating surviving stress fibers in cells. (g) Box-plot of total actin fiber lengths extracted from fluorescence images for cells with vimentin intermediate filaments and cells with vimentin-Y117L droplets with and without Cytochalasin B treatment.

To further explore the consequences of the wetting of vimentin droplets on actin fibers, we treat the cells with Cytochalasin B, which binds to the polymerizing end of actin filaments and prevents elongation^40^. However, the depolymerization step of actin treadmilling remains unaffected and thus the actin filaments eventually dissolve. When we treat cells containing normal network of VIFs with 5 *μ*g/ml of Cytochalasin B for 1 hour, they lose almost all actin fibers, as can be seen from the lack of fibers in the actin image (magenta) of cells shown in Fig. 6f. By contrast, cells with vimentin-Y117L droplets retain a substantial amount of actin fibers after the same drug treatment, as seen in Fig. 6f. Cells containing VIFs and cells containing vimentin-Y117L droplets have an average of ∼731 *μ*m and ∼782 of actin fibers per cell respectively before drug treatment. In drug treated, VIF-containing cells the total fiber length is almost 5-fold lower than the control population. In contrast, vimentin-Y117L cells have half the length of actin fibers even after drug treatment. This can be seen by comparing the sharp decrease between first and second boxes with the less drastic decrease between third and fourth boxes in Fig. 6g.

Microscope images show that the actin fibers are partially coated by distinct droplets, as can be seen by the fluorescence image of vimentin droplets (green) and actin fibers (magenta) shown in actin Fig. 6b. Therefore, we expect that only those coated sections of the fibers to be protected after Cytochalasin B treatment. Surprisingly however, some drug-treated cells containing vimentin-Y117L exhibit continuous actin fibers, as can be seen in the image of the actin shown in magenta in Fig. 6f. One possibility to account for this behavior is that a thin layer of vimentin, which cannot be imaged with a fluorescence microscope, coats the entirety of these fibers.

### Wild type vimentin forms liquid droplets which colocalize with actin fibers

The results regarding LLPS and wetting, though conclusive, are from vimentin-Y117L droplets which cannot form VIFs. Therefore, we explore the properties of wild type vimentin, capable of forming filaments, during assembly. We treat MEF cells expressing m-Emerald tagged VIFs with 20% (v/v) diluted culture medium which reversibly dissolves VIFs^41,42^. When we replace the diluted medium with fresh complete medium, the vimentin reassembles in a series of steps beginning with non-filamentous punctate structures distributed throughout the cell, as shown in Fig. 7a. When two of these puncta come in contact with one another, they can merge, as seen inside the yellow circle in Figs. 7b-d, confirming that the puncta are liquid droplets likely formed through LLPS. These droplets eventually form VIFs; thus, LLPS appears to play a role in the assembly of VIFs.

**Figure 7.**
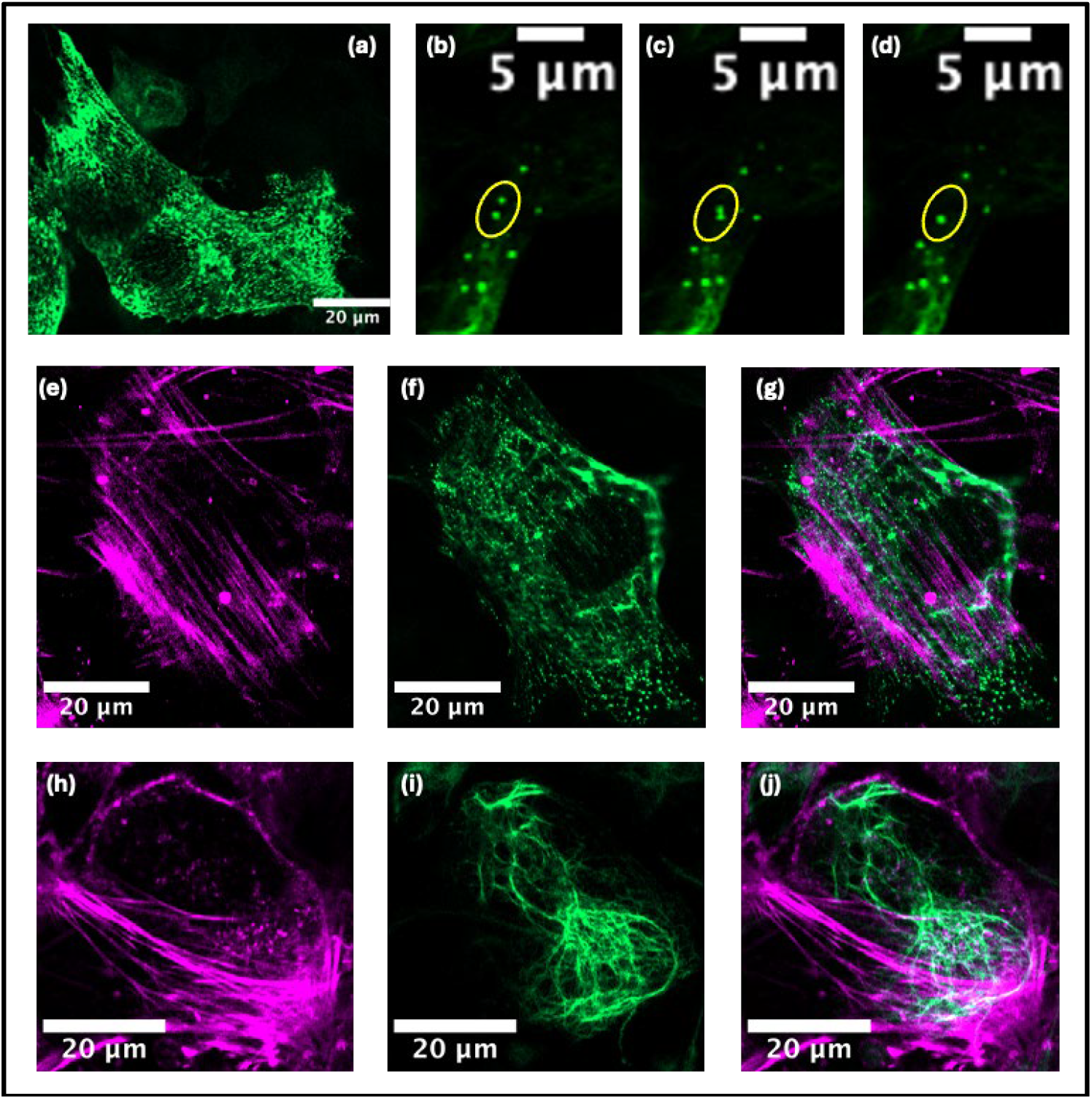
Vimentin filament precursors are liquid droplets and colocalize with actin fibers. (a) Vimentin droplets (green) formed as a precursor to filaments. (b)-(d) Two Vimentin droplets merging inside the yellow oval. (e),(f) Actin fibers (magenta) and Vimentin filament precursor droplets (green) of a cell. (g) Overlay image showing the droplets colocalize with actin fibers well. (h),(i) Actin fibers (magenta) and Vimentin filaments (green) of a cell. (j) Overlay image showing very little colocalization between the two networks.

To explore the behavior of the precursor droplets, we observe how they interact with actin fibers. These droplets colocalize with actin fibers, which can be seen from the image of actin network (magenta) and the vimentin droplets (green), as well as the overlaid image shown in Figs. 7e, f and g respectively. In the same cells, when vimentin exists as filaments,they show no obvious overlap with actin fibers, as can be seen by comparing the actin network (magenta) and the vimentin network (green), as well as the overlaid image shown in Figs. 7h, i and j respectively. The droplets differ from VIFs only in their physical state, thus the enhanced overlap between vimentin droplets and actin fibers indicate that actin-vimentin interaction is governed by vimentin’s state.

## Conclusions

In this paper, we show that vimentin-Y117L forms liquid condensates through phase separation in MEF cells. The exact composition of these droplets is unknown; they must be composed of the essential sub-units that ultimately can form whatever structure vimentin produces. The vimentin-Y117L mutant cannot elongate beyond tetramers and ULFs, and therefore these sub-units must be sufficient to form the condensates^15,30^. The phase separation of proteins is strongly linked to intrinsically disordered regions in their structure, which can lead to weak, non-specific attractions that drive the phase separation^27,28^. Vimentin-Y117L has IDRs on both its N- and C-terminals, and thus its phase separation is not surprising. Interestingly, wild type vimentin has the same IDRs as vimentin-Y117L and therefore should exhibit similar phase behavior. Indeed, we observe that wild type vimentin forms liquid droplets. This suggests that these droplets may be phase separated condensates.

In this paper, we show that vimentin intermediate filaments nucleate and grow out of these droplets during assembly. This implies that these condensates are important for VIF assembly. Moreover, wild type vimentin cannot form filaments when its N-terminal IDR is removed^43^; this suggests that weak interaction driven by IDRs may also contribute to VIF assembly^44^.

This VIF network is one of three filamentous protein networks in the cell, filamentous actin and microtubules being the other two. The growth of each of these filaments requires addition of sub-units with both the correct position and correct orientation, which will slow the extension of the filaments. Increasing the local concentration at the growing tip of the filament will increase the growth rate. Interestingly, this seems to be accomplished by phase separated condensates in each case. For actin filaments, polymerization occurs 14 times faster inside N-WASP condensates^10^. For microtubules, assembly occurs in centrosomes, which are phase separated condensates formed by its auxiliary proteins^11^. Our results suggest that a similar condensate-mediated process may be involved in VIF assembly; vimentin sub-units are concentrated inside droplets where filaments can nucleate and grow. Interestingly, vimentin can form these condensates by itself, unlike actin or microtubules. However, we do not know exactly which vimentin sub-units make up the wild type condensates. The attractive interaction due to IDRs are essential for phase separation to form condensates and in turn increase the concentration during the filament growth process; these condensates are implicated in the formation of the filaments themselves. It is therefore conceivable that the formation of VIFs may depend on the same attractive interaction due to the IDRs.

In this paper, we show that vimentin-Y117L condensates colocalize extensively with actin fibers and remain attached to them. This interaction is driven by wetting. It is strong enough to coat the fibers continuously and prevent their disassembly under Cytochalasin B treatment; such a strong wetting interaction is rare in proteins. We similarly observe wild type droplets colocalizing on actin fibers before assembling into filaments. This suggests that actin plays a role in the assembly of VIFs. Actin fibers may act as a surface on which vimentin droplets nucleate. Additionally, being colocalized on actin may help the orient the vimentin sub-units to better incorporate into the growing filaments. Vimentin filaments growing out of these droplets may use actin fibers as their template; as these VIFs elongate, they eventually detach from the actin fibers. This might be due to a mismatch in their persistence lengths. Vimentin intermediate filaments have a lower persistence length^45–47^ and thus must be straightened to stay attached to the actin fibers. This becomes energetically unfavorable with increasing length of the VIF, causing them to detach. This demonstrates how components of the cytoskeletal network may influence one another’s assembly through phase separation and wetting.

## Materials and Methods

### Cell culture and live actin staining

Mouse Embryonic Fibroblast vimentin-mEmerald (MEF vim-mEmerald), Mouse Embryonic Fibroblast vimentin-Y117L (MEF vim-Y117L) and Mouse Embryonic Fibroblast vimentin-Y117L-mEmerald (MEF vim-Y117L-mEmerald) cell lines were gifts from the lab of Prof. Robert Goldman of Northwestern University. MEF cells are cultured in Dulbecco’s modified Eagle’s Medium (DMEM, Corning, catalog no. 10-013-CV) supplemented with 10% fetal bovine serum (FBS, Avantor, catalog no. 97068-085) and 1% Penicillin/Streptomycin (Corning, catalog no. 30-002-CI). For live actin staining, cells are cultured for at least 24 hours and then treated with 100nM SiR-actin (Cytoskeleton Inc., catalog no. CY-SC001) for 12 hours. After treatment, the cells are washed with 1X Phosphate buffered saline (PBS, Corning, catalog no. 21-040-CV) and treated with fresh complete culture medium before imaging.

### Immunostaining

Cells are fixed with 4% paraformaldehyde, permeabilized with 0.1% Triton-X-100 (Thermo Fisher Scientific, catalog no. A16046.AE), blocked with blocking buffer^1^, treated with primary antibody overnight at 4^o^C and treated with secondary antibody and phalloidin (Abcam, ab176753) for 1 hour at room temperature. Next, they were treated with DAPI at room temperature for 10 minutes, washed and imaged. For details of primary and secondary antibodies, refer to Table 1. ^1^blocking buffer: 22.52 mg/ml glycine (Aldrich, catalog no. 241261), 1% bovine serum albumin (BSA, Sigma-Aldrich, catalog no. A3733) and 1% Tween-20 (Sigma-Aldrich, catalog no. P9416) in 1X PBS buffer.

**Table 1.** Specific primary and secondary antibodies, their catalog numbers and concentrations

| Antibody | Primary/<br>Secondary | Company, Catalog No. | Dilution |
| --- | --- | --- | --- |
| Anti-vimentin rabbit polyclonal | Primary | Abcam, ab45939 | 1:1000 |
| Anti-paxillin mouse monoclonal | Primary | Thermo Fisher Scientific, AHO0492 | 1:100 |
| Goat anti-rabbit IgG Alexa Fluor 647 | Secondary | Thermo Fisher Scientific, A21244 | 1:500 |
| Goat anti-mouse IgG Alexa Fluor 555 | Secondary | Thermo Fisher Scientific, A28180 | 1:500 |

### 1,6-Hexanediol treatment

MEF cells are cultured for at least 24 hours and stained with SiR-actin. They are treated with 0.5% (w/v) 1,6-Hexanediol (Sigma-Aldrich, 240117) in complete culture medium for 2 minutes, fixed with 4% paraformaldehyde and imaged. For immunostaining, MEF cells are cultured for at least 24 hours, treated with 0.5% (w/v) 1,6-Hexanediol in complete culture medium for 2 minutes, fixed with 4% paraformaldehyde and stained for vimentin, paxillin and actin before imaging.

### Diluted media treatment

MEF cells are grown for at least 24 hours and treated with SiR-actin. While the cells are on the microscope, the media is aspirated leaving some liquid behind to prevent cells from drying out. Immediately, the 20% (v/v) diluted culture media in water is added and imaging is continued. For recovery, the diluted media is aspirated, and fresh complete culture media is added to the cells.

### Colocalization threshold calculation

Colocalization between actin and vimentin images are calculated by using the “colocalization threshold” plugin in Fiji. It follows the Costes method for determining thresholds; if the intensities in the two channels are above the threshold values of the respective channels for a particular pixel, then it is considered colocalized. The two thresholds are determined iteratively to ensure that the Pearson’s correlation coefficient for all non-colocalized pixels is zero and the coefficient is greater than zero for all colocalized pixels.

### Cytochalasin B treatment

MEF cells are grown for at least 24 hours and treated with 5*μ*g/ml of Cytochalasin B (Millipore-Sigma, catalog no. 250225) in complete culture medium for 1 hour and fixed with 4% paraformaldehyde. Cells are stained with Phalloidin for actin and DAPI (Thermo Fisher Scientific, catalog no. 62247) for the nucleus before imaging.

### Droplet tracking and motion analysis

Droplets are tracked from fluorescence microscopy images using the “trackmate” plugin in Fiji. First, the particles are detected using a Laplacian of Gaussian (LoG) detector for every single frame and quality threshold is adjusted to ensure reliable detection. The tracking of particles is done by “advanced Kalman tracking”. The X,Y and time coordinates of the particles are exported as .xml files and the following analyses are performed using Matlab.

We calculate the mean squared displacement (⟨Δ*r*^2^⟩) for all possible time differences (Δ*t*) in the trajectories and plot them on a log-log scale. For diffusive motion, the governing equation of motionis ⟨Δ*r*^2^⟩ = 4*D \** Δ*t*, where D is the diffusion coefficient of the droplets. Thus, on log-scale, the equation becomes *log*(⟨Δ*r*^2^⟩) = *log*(4*D*) + *log*(Δ*t*), which has a slope of 1. By fitting the data to a linear regression model, we can estimate the intercept fo the plot from which the diffusion coefficient (D) can be calculated.

For ballistic, or directional, motion the governing equation of motion is Δ*r* = *v* * Δ*t*, where v is the velocity of the particle. So, the root mean square displacement is related to time as ⟨Δ*r*^2^⟩ = *v*^2^ *\** Δ*t*^2^. On a log-scale, the equation becomes *log*(⟨Δ*r*^2^⟩) = 2 * *log*(*v*) + 2 * *log*(Δ*t*), which results is a slope of 2. By fitting the ballistic part of the data to a linear regression model, we can estimate the intercept of the plot from which the directional velocity of the droplets can be calculated. The codes for the Matlab analyses are provided on GitHub.

### Actin fiber detection

The actin network image of the cells are pre-processed by enhancing their brightness and contrast for better identification whenever necessary. The detection of the fibers is done by an original Python based algorithm using the computer vision library OpenCV2^48^.

A Gaussian blur is applied on the image to remove high frequency noise. Then a Canny edge detection identifies the filament boundaries; this works by examining the gradient of the image at each pixel in all directions. Pixels that are local gradient maxima along their gradient direction are edge candidates. Then we set an upper and a lower threshold for gradients. All edge candidates above the upper threshold are considered true edges; all edge candidates below the lower threshold are rejected. Whether edge candidates between the two thresholds are rejected or not depends on their spatial position. If they are contiguously connected to other true edges, then they are considered true edges themselves, if not they are rejected. The detected edges are skeletonized whenever necessary to a width of 1 pixel.

The gradient direction of an edge is always perpendicular to the filament itself, and for a filament, there are two edge pixels corresponding to the opposite sides of the filament. We detect the filaments by the existence of two such edges that are within 0.7 *μ*m along the gradient direction of the original, unblurred image; this threshold is set for individual images. A “filament pixel” is placed on the position between two edge pixels. The filament pixels are connected together by drawing a circle of fixed radius around them and then skeletonizing. Filaments with an in-image area smaller than a specific threshold are eliminated. We estimate the length of individual filaments by contourizing them with OpenCV2, which is done by calculating their perimeter and dividing by two. This also reduces the length impact of spurious pixels that might emerge from skeletonization. From individual filament lengths, we obtain the total filament length in cells. All the relevant codes are provided on GitHub.

