## Supplementary Figures for "Phase separated liquid vimentin droplets stabilize actin fibers through wetting"

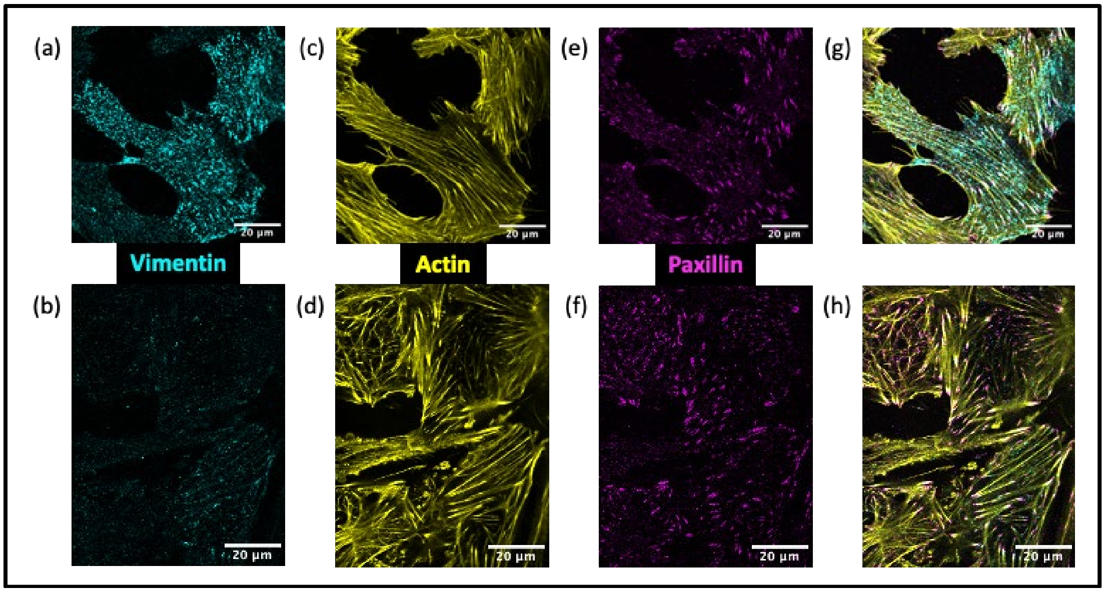


**Supplementary Figure 1| Effect of 1,6-Hexanediol on MEF cells.** (a),(b) Vimentin-Y117L droplets (cyan) before and after 0.5% 1,6-Hexanediol treatment for 2 minutes. (c),(d) Actin fibers (yellow) before and after 0.5% 1,6-Hexanediol treatment for 2 minutes. (e),(f) Paxillin (magenta) staining before and after 0.5% 1,6-Hexanediol treatment for 2 minutes. (g),(h) Overlay image of vimentin-Y117L (cyan), actin (yellow) and paxillin (magenta) staining before and after 0.5% 1,6-Hexanediol treatment for 2 minutes.

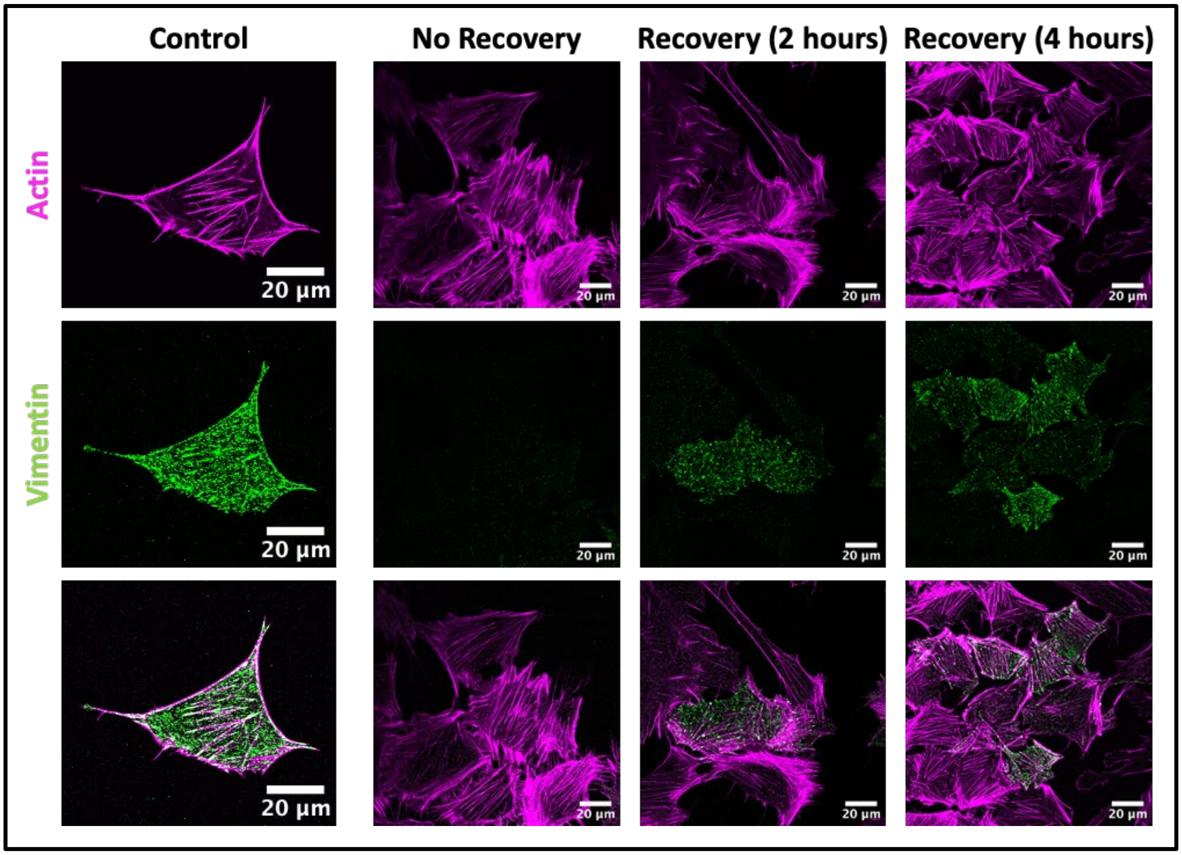


**Supplementary Figure 2| Recovery of vimentin-Y117L droplets in complete culture medium after 0.5% 1,6-Hexanediol treatment for 2 minutes.**
